# Whole-genome sequencing analysis of copy number variation (CNV) using low-coverage and paired-end strategies is efficient and outperforms array-based CNV analysis

**DOI:** 10.1101/192310

**Authors:** Bo Zhou, Steve S. Ho, Xianglong Zhang, Reenal Pattni, Rajini R. Haraksingh, Alexander E. Urban

## Abstract

**Background:** CNV analysis is an integral component to the study of human genomes in both research and clinical settings. Array-based CNV analysis is the current first-tier approach in clinical cytogenetics. Decreasing costs in high-throughput sequencing and cloud computing have opened doors for the development of sequencing-based CNV analysis pipelines with fast turnaround times. We carry out a systematic and quantitative comparative analysis for several low-coverage whole-genome sequencing (WGS) strategies to detect CNV in the human genome.

**Methods:** We compared the CNV detection capabilities of WGS strategies (short-insert, 3kb-, and 5kb-insert mate-pair) each at 1x, 3x, and 5x coverages relative to each other and to 17 currently used high-density oligonucleotide arrays. For benchmarking, we used a set of Gold Standard (GS) CNVs generated for the 1000-Genomes-Project CEU subject NA12878.

**Results:** Overall, low-coverage WGS strategies detect drastically more GS CNVs compared to arrays and are accompanied with smaller percentages of CNV calls without validation. Furthermore, we show that WGS (at ≥1x coverage) is able to detect all seven GS deletion-CNVs >100 kb in NA12878 whereas only one is detected by most arrays. Lastly, we show that the much larger 15 Mbp Cri-du-chat deletion can be readily detected with short-insert paired-end WGS at even just 1x coverage.

**Conclusions:** CNV analysis using low-coverage WGS is efficient and outperforms the array-based analysis that is currently used for clinical cytogenetics.

## INTRODUCTION

A large portion of human genetic diversity is contributed by CNVs [1–5]. Many CNVs, typically small deletions or duplications, are common, i.e. present at an overall frequency of >1% in the human population [3–6]. Large CNVs are relatively rare and are often associated with human disease [7–14]. Having technologies available for the reliable and accurate detection and characterization of CNVs in a given human genome is highly relevant for both clinical diagnostics and basic research. Microarray-based CNV analysis has become a first-tier clinical cytogenetics procedure in patients with unexplained developmental delay/intellectual disability [15], autism spectrum disorder [16], multiple congenital anomalies [17], and cancer [13, 14].

The highest sensitivity and resolution in CNV detection is achieved through deep-coverage, paired-end whole-genome sequencing (WGS) [5]. However, the cost for what is currently the standard for deep-coverage WGS (>30x coverage using short-insert paired-end reads) is still considerably higher than for that of arrays; turnaround time is much longer since the samples have to go through an offsite core, and the computational requirements are also very substantial regarding hardware and time. Analysis by deep-coverage WGS methods can not only detect CNVs but also SNPs, short insertions and deletions as well as, with some limitations, sequence variants that are quite challenging to parse out such as inversions and retrotransposition events. However, for clinical cytogenetic applications such types of variants are for the most part not yet interpretable as to their effects.

With the advent of bench-top high-throughput DNA sequencers it is now possible to perform low-coverage WGS on-site instead of through a sequencing core facility. To make the most beneficial use of this option, i.e. to control per-sample-costs as well as turnaround times, it seems beneficial to use a strategy of lower sequencing coverage (i.e. 1x-5x genomic coverage) with sample multiplexing to be cost-effective while carefully weighing the options of short-insert versus long-insert paired-end (i.e. mate-pair) library preparation.

While other recent studies have demonstrated that WGS, including low-coverage WGS, is effective for CNV detection in clinical samples [18–20], systematic, quantitative, and direct performance comparisons for CNV analysis between various low-coverage WGS strategies, against deep-coverage WGS, and against arrays, are needed to fully assess the feasibility of replacing arrays with low-coverage WGS and to guide researchers in their choices for specific settings. Here, we compared the CNV-detection performances of several low-coverage WGS strategies against each other and also against commercially available arrays. We performed CNV analysis in the genome of the 1000-Genome-Project CEU subject NA12878 (probably the best studied genome to date [3, 21, 22]) using standard 350 bp short-insert WGS, 3kb-insert mate-pair WGS, and 5kb-insert mate-pair WGS, each at 1x, 3x, and 5x coverages. For benchmarking, we used a Gold Standard (GS) set of validated CNVs for NA12878 and determined the number of GS CNVs detected by each low-coverage WGS strategy. The GS set contains only high confidence CNVs derived from the 1000 Genomes Project and supported by multiple orthogonal methods [23, 24]. This approach was also used in Haraksingh *et al* [24] for the benchmarking of CNV detection in NA12878 from 17 commercially available arrays and thus allows for the performance comparison between low-coverage WGS and the arrays from Haraksingh *et al* [24] to be conducted in a direct and unbiased manner.

## METHODS

Sample library construction, sequencing, alignment, CNV analysis, array processing, and NCBI accession numbers are described in Supplementary Materials and Methods.

## RESULTS

### CNV detection in WGS

From short-insert and mate-pair WGS of NA12878, we performed CNV detection using both read-depth and discordant read-pair analysis (Figure 1a). For read-depth analysis, we used CNVnator [25] with 5kb bin size. Discordant read-pair analysis was performed using LUMPY [26] with segmental duplications excluded from the analysis. CNVs that overlap problematic regions such as reference gaps, the MHC cluster, and ENCODE blacklist regions [27] were filtered out (see Methods). Afterwards, the union of CNV calls from both analyses was used as the final call set for benchmarking using the GS CNVs as well as comparison with the array calls [24]. At 1x, 3x, and 5x coverages, short-insert WGS detects 182, 405, and 535 autosomal CNVs respectively; 3kb mate-pair WGS detects 452, 689, and 747 respectively; 5kb mate-pair WGS detects 496, 571, and 725 respectively (Supplementary Table S1, Figure 1b).

**Figure 1.**
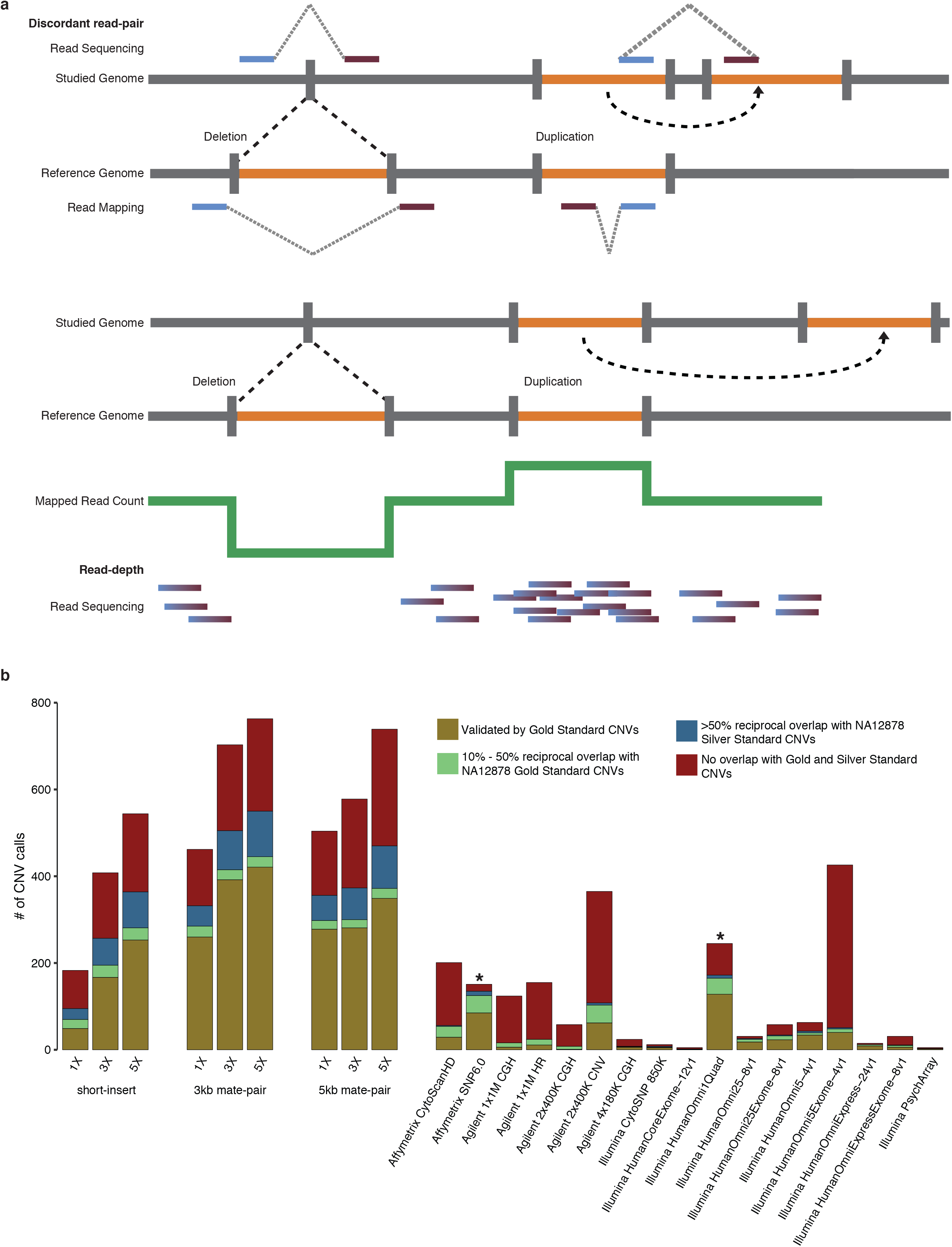
Comparisons of CNV Calls by WGS and Arrays. (**a**) Schematic diagram of detection of CNVs (deletions and duplications) using discordant read-pair and read-depth analysis [1, 25, 39]. Using discordant read-pair analysis, deletions are detected when the distance of alignment to the reference genome between read pairs are closer than the expected insert size of the library, and duplications are detected when the orientation of the aligned read pairs are inversed. Using read-depth analysis, deletions and duplications are detected when there is a pronounced decrease and increase, respectively, in alignments of reads spanning a genomic region relative to the average number of alignments over the genome. (**b**) Numbers of autosomal CNVs in the genome of subject NA12878 called from short-insert, 3kb mate-pair, and 5kb mate-pair libraries sequenced at 1X, 3X, and 5X coverage compared against previous array calls [24, 28]. Array-based CNV calls were made according to platform-specific algorithms [24], and WGS-CNV calls were made by combining discordant read-pair and read-depth analysis. Gold: autosomal CNVs (overlap ≥50% reciprocally with NA12878 GS CNVs). Green: 10%-50% reciprocal overlap with NA12878 GS CNVs. Blue: <10% reciprocal overlap with GS CNVs, ≥50% reciprocal overlap with NA12878 Silver Standard CNVs. Red: no overlap (<10% overlap with GS CNVs and <50% overlap with Silver Standard CNVs). Benchmarking was performed taking CNV type into account with the exception of Affymetrix SNP 6.0^*^ [28] and Illumina HumanOmni1Quad^*^ [24] where CNV type information was not available.

### CNV-detection performance comparison

We obtained the NA12878 CNV calls by each of 17 currently commercially available high-density oligonucleotide arrays from Haraksingh *et al* [24]. These arrays represent three different technologies: array CGH (aCGH) from Agilent (*n*=5), SNP genotyping arrays from Illumina (*n*=10), and aCGH/SNP combination arrays from Affymetrix (*n*=2). Two technical replicates had been performed for each array hybridization, and CNVs were called using both array platform-specific software as well as platform-agnostic software Nexus from Biodiscovery except for Affymetrix SNP 6.0 where the platform-specific calls (one replicate available) were obtained from an earlier study [28].

We benchmarked the CNV calls from short-insert and mate-pair WGS using the same approach as described in Haraksingh *et al* [24], where the capabilities of various array platforms were assessed by the numbers of detected CNVs in the NA12878 genome that reciprocally overlap a GS set of NA12878 CNVs. GS CNVs were compiled from 8x-coverage population-scale sequencing (data available on 1000genomes.org) and analysis of 2,504 individual genomes [23]. They are of high-confidence and supported by multiple lines of evidence that include PCR confirmation, aCGH, and discovery from multiple CNV analysis tools. The false-positive rate is estimated to be very low (3.1%) [24]. The CNVs in this GS set range from 50 bp to 453,312 bp with 1,941 and 135 autosomal deletions and duplications respectively [24]. While most GS duplications are >10 kb, all CNVs <1kb are deletions. Seven deletions are >100 kb (Table 1). A NA12878 Silver Standard set of CNVs was also used for benchmarking (also obtained from Haraksingh *et al* [24]) which consists of CNVs called using only CNVnator [25] from 60x-coverage 2×250 bp short-insert sequencing data from the 1000 Genomes Project. For our analysis, we filtered out Silver Standard CNVs that overlapped reference gaps [29, 30] by >50% (Supplementary Figure S1).

**Table 1.** Detection of Gold Standard deletions > 100 kb

| Gold Standard Deletion | Size (bp) | WGS Detection | Array Detection (Haraksingh *et al* 2017) |
| --- | --- | --- | --- |
| chr3:162,514,471-162,625,647 | 111,176 | All WGS | All Agilent arrays<sup>a</sup> |
| chr4:69,375,591-69,491,543 | 115,952 | All WGS | Affymetrix Cytoscan HD<br>Affymetrix SNP 6.0<br>Agilent 2x400K CNV<sup>a</sup><br>Illumina HumanOmni1Quad-v1<br>Illumina HumanOmni2.5-8v1<br>Illumina HumanOmni25Exome-8v1<br>Illumina HumanOmni5-4v1<br>Illumina HumanOmni5Exome-v1 |
| chr4:70,122,981-70,231,746<sup>b</sup> | 108,765 | All WGS | Affymetrix SNP 6.0<br>Agilent 2x400K CNV<sup>a</sup><br>Illumina HumanOmni1Quad-v1<br>Illumina HumanOmni2.5-8v1<br>Illumina HumanOmni25Exome-8v1 |
| chr6:78,892,808-79,053,430 | 160,622 | All WGS | Affymetrix Cytoscan HD<br>Affymetrix SNP 6.0<br>Agilent 2x400K CNV<sup>a</sup><br>Illumina HumanOmni1Quad-v1<br>Illumina HumanOmni2.5-8v1<br>Illumina HumanOmni25Exome-8v1<br>Illumina HumanOmni5-4v1<br>Illumina HumanOmni5Exome-v1<br>Illumina HumanOmniExpressExome 1.2 |
| chr8:39,195,825-39,389,230 | 193,405 | All WGS | Affymetrix Cytoscan HD<br>Affymetrix SNP 6.0<br>Agilent 2x400K CNV<sup>a</sup><br>Illumina HumanOmni1Quad-v1<br>Illumina HumanOmni2.5-8v1 |
| chr11:4,248,265-4,353,229 | 104,964 | All WGS | Affymetrix SNP 6.0<br>Agilent 2X400K CNV<br>Agilent 1X1M HR |
| chr19:20,595,835-20,717,950 | 122,115 | All WGS | All arrays except Illumina Psych Array |

<sup>a</sup>called as high copy duplication

As was done for array CNV calls [24], the CNV calls from each WGS library at each sequencing coverage were benchmarked by first determining the number of CNVs with boundaries overlapping those of a GS CNV by ≥50% reciprocally and the number of CNVs that overlap of a GS CNV ≥10% reciprocally but <50%. From the CNVs that do not overlap a GS CNV by ≥10% reciprocally, we next determined the number that overlap a Silver Standard CNV by ≥50% reciprocally (Supplementary Table S1, Figure 1b). For array data [24], because more CNV calls and GS-CNV overlaps resulted from the platform-specific CNV analysis overall, we chose to use the array platform-specific calls for comparison. Moreover, because the results in the two technical replicates for each array platform did not show significant differences and only one replicate was available for the Affymetrix SNP 6.0 platform-specific analysis [24], we chose to use CNV calls from the first replicate of each platform. We emphasize that we benchmarked all WGS and array CNV calls by taking the type of CNV (deletion or duplication) into account which was not done in Haraksingh *et al* [24].

With the exception of short-insert WGS at 1x coverage, WGS detects drastically more CNVs and GS CNVs than any of the arrays (Figure 1b). CNV calls from WGS are also accompanied by a smaller percentages of calls without validation (i.e. CNV calls that are not of high confidence). Validation is hereinafter defined as overlap by a minimum of 10% reciprocally with a GS CNV or >50% reciprocally with a Silver Standard CNV. At coverages 1x, 3x, and 5x and by >50% reciprocal overlap, short-insert WGS detected 48, 164, and 244 GS autosomal CNVs respectively; 3kb mate-pair WGS detected 250, 378, and 405 GS autosomal CNVs respectively; and 5kb mate-pair WGS detected 270, 274, and 335 GS autosomal CNVs respectively (Supplementary Table S1).

As expected, read-depth analysis resulted in higher rates of CNV detection with increasing coverage but showed similar numbers of CNV calls across different WGS libraries (Supplementary Figure S2a, Supplementary Table S2). Discordant read-pair analysis resulted in consistently more CNV calls than read-depth analysis for mate-pair libraries at all coverages and for short-insert library at 5x coverage (Supplementary Figure S2b, Supplementary Table S3). Generally, in read-depth analysis, the greater the fraction of uniquely-mapped supporting reads, the higher the confidence in the CNV call. Filtering based on this parameter can be done through the *q0* value reported by CNVnator [25] (Supplementary Table S2). In Abyzov *et al* [25], the *q0* threshold was set as 0.50 indicating that CNVs supported by >50% of reads with a mapping quality of zero were filtered out. To understand how such filtering affects our read-depth analysis, we benchmarked filtered (*q0* threshold = 0.50) and unfiltered CNV calls (Supplementary Figure S2a,c). We find that the overall number of GS CNVs detected did not noticeably change with the *q0* filter. However, number of Silver Standards detected dramatically decreases (Supplementary Figure S2c). In addition, the number of non-validated calls decrease less dramatically but very substantially nonetheless (~12%-30%), consistent with the average decrease in false discovery-rate for samples studied in Abyzov *et al*[25].

Of the arrays, Illumina HumanOmni1 Quad (now discontinued) detects the most GS CNVs (165) [24]; however, even at 1x coverage, 3kb- and 5kb-mate-pair WGS detects almost twice as many GS CNVs (275 and 290 respectively) (Supplementary Table S1, Figure 1b). While Agilent 2x400K CNV and Illumina HumanOmni5Exome-4v1 detect comparable numbers of CNV to that of short-insert WGS at 3x coverage and to that of mate-pair WGS at 1x coverage, the vast majority of CNVs detected in these two arrays are not validated (Figure 1b, Supplementary Figures S3). This is in contrast to low-coverage WGS results where the majority of CNVs detected for all libraries and at all sequencing coverages have validation (Figure 1b, Supplementary Figure S3, Supplementary Table S1).

WGS (3kb-mate-pair) at 5x coverage results in the most number of GS CNVs detected (429) and also the lowest percentage without validation (28.5%) (Figure 1b, Supplementary Figure S3, Supplementary Table S1). The percentages of CNV calls without validation range from 28.5%-37.1% for mate-pair WGS and 33.6%-48.4% for short-insert WGS (Supplementary Figure S4, Supplementary Table S1). These percentages for WGS are smaller than those for most arrays except Illumina HumanOmniExpress and HumanOmni25 arrays (Supplementary Figure S3). These two arrays, however, made a very low number calls (15 and 31 respectively) compared to WGS (Figure 1b, Supplementary Table S1). Affymetrix SNP 6.0 also has a higher overall validation rate than that of WGS, but information for whether a CNV call is a deletion or duplication is not available for this array dataset [24, 28]. It is uncertain how its validation rate will change if this information can be taken into account (as for the WGS and other array datasets).

When deletions and duplications are analyzed separately, the validation rates for deletions are higher than for duplications (56.1%-75.6% vs 34.8%-49.4%) in WGS whereas arrays show a much wider variability (Supplementary Figure S4, Supplementary Table S1). For deletions, the Agilent CGH arrays, Affymetrix CytoScanHD and the Illumina CytoSNP, HumanCoreExome, HumanOmniExpressExome-8v1, and Psych arrays have validation rates between 15% to 50%. The Illumina HumanOmni arrays (except HumanOmniExpressExome-8v1) have validation rates between 54.9%-83.3%, but none of these arrays detected more than 70 deletions. For duplications, the Agilent CGH, HumanOmni5Exome, and Affymetrix CytoScanHD have validation rates between <1% to 20%. All other arrays detected no more than 7 duplications. The Illumina CytoSNP, HumanCoreExome, HumanOmni25-8v1, and Psych arrays detected 4, 1, 5, and 2 duplications respectively, and all were validated (Supplementary Table S1). The HumanOmni arrays (except HumanOmni5Exome) have validation rates between 50.0%-85.7% detecting duplications ranging from 2 to 7. For both deletions and duplications, the numbers of Gold and Silver Standard CNVs detected and the overall validation rates are much higher for WGS compared to the arrays (Supplementary Figure S4).

### Sensitivity of GS CNV detection

Overall, low-coverage WGS is able to detect on average ~5-fold more GS CNVs (>1 kb) compared to arrays by ≥50% reciprocal overlap (Figure 2a). For example, the most sensitive array (as measured by the number of GS CNVs detected), Illumina HumanOmni1 Quad (now discontinued), detects less than one third as many GS CNVs than the most sensitive WGS method (3kb-mate-pair at 5x coverage). The second least sensitive WGS method (short-insert at 3x coverage) is still more sensitive than Illumina HumanOmni1 Quad. Even the least sensitive WGS method (short-insert at 1x coverage) is more sensitive than 14 out of the 17 arrays. Moreover, mate-pair WGS is able to detect >50% of GS CNVs in the 5kb-10kb size range (Figure 2b-d).

**Figure 2.**
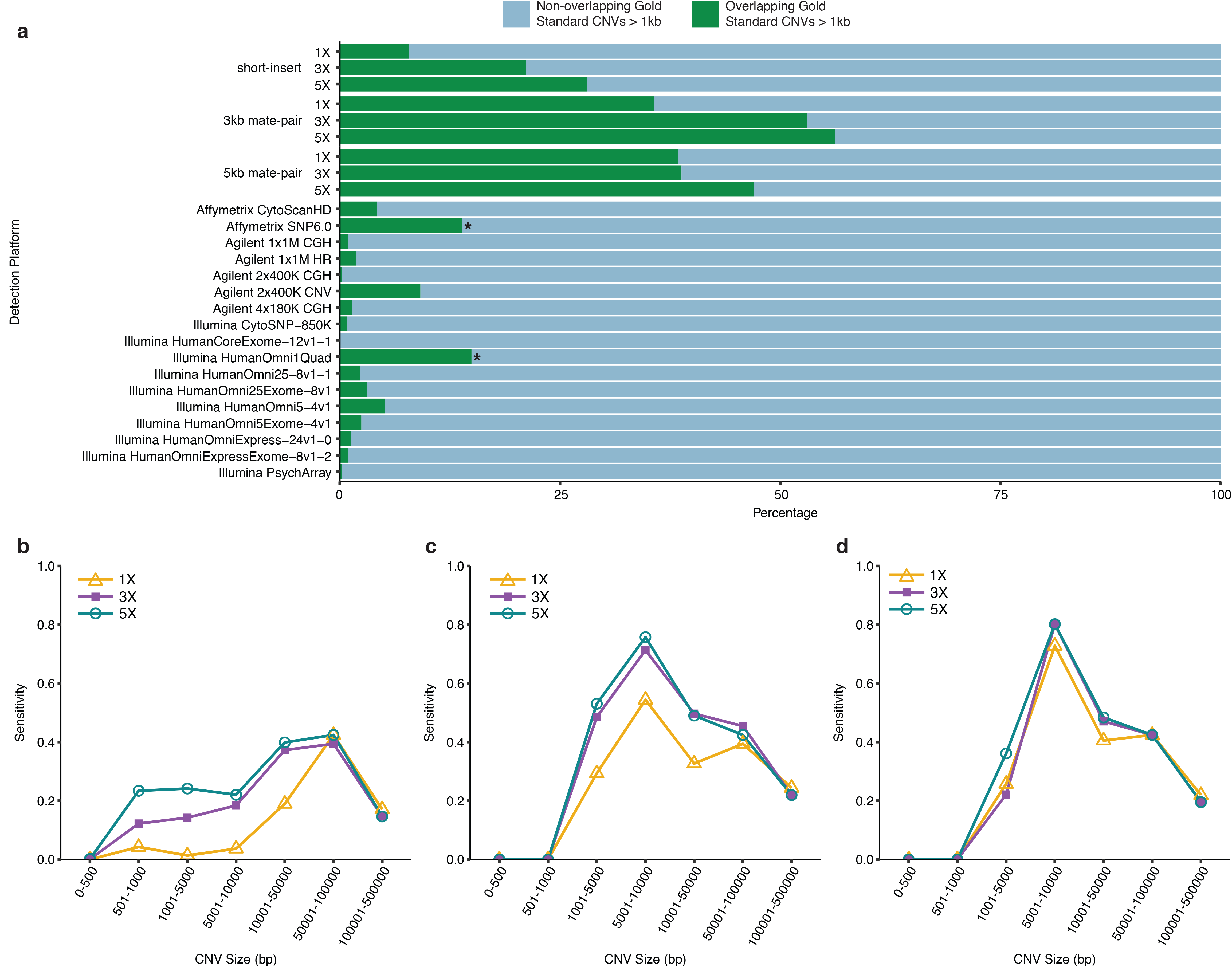
Sensitivity of WGS Detection of NA12878 GS CNVs (>1 kb). (**a**) Sensitivity of (>1 kb) GS CNV detection across WGS libraries and array platforms as determined by the ratio of detected autosomal GS CNVs to total number of autosomal GS CNVS. Array-based CNV calls were made according to platform-specific algorithms, and WGS CNV calls were made by combining discordant read-pair and read-depth analysis. Green: autosomal CNVs that overlap >50% reciprocally with NA12878 GS CNVs. Blue: total number of NA12878 GS autosomal CNVs. Sensitivity of GS CNV detection in different size ranges from (**b**) short-insert, (**c**) 3kb mate-pair, and (**d**) 5kb mate-pair libraries at sequencing coverages 1X, 3X, and 5X. CNVs were called by combining discordant read-pair and read-depth analysis.

While CNV detection increases with additional coverage for all WGS libraries, the increase is non-linear. The highest increases are from 1x to 3x coverage for short-insert and 3kb-mate-pair WGS (Figure 1b). While more CNVs are consistently detected from mate-pair WGS compared to short-insert WGS, interestingly, more total CNVs and GS CNVs are detected from 3kb-mate-pair WGS than from 5kb-mate-pair WGS at 3x and 5x coverages, respectively (Figure 1b, Figure 2a). In addition, while additional coverage is associated with overall increases in the detection of GS CNVs, this increase is less obvious as CNV sizes increase to >50kb (Figure 2b-d). In short-insert WGS, with additional coverage, the most drastic gains in the GS CNVs detected are between 5kb to 50kb (Figure 2b). This is similar for mate-pair WGS though less pronounced (Figure 2c, d).

### Size distribution of CNV calls

The sizes of NA12878 CNVs detected from short-insert and mate-pair WGS range from 100bp to 500kb and 1kb to 500kb respectively (Figure 3a-c, Supplementary Tables S2-S4). Read-depth and discordant read-pair analysis detect CNVs in different size ranges (Figure 3d-i, Supplementary Figure S5). Overall, WGS detects CNVs in a wider size distribution compared to arrays (Supplementary Figure S5). Since bin size was set to 5kb (see Methods), all resulting CNVs detected are ≥5 kb from read-depth analysis (Figure 3d,e,f). As expected, the size distributions of CNVs called by read-depth analysis are very similar (Figure 3d-f) for the various WGS libraries, whereas discordant read-pair analysis shows more variability (Figure 3g-i) reflecting the different insert sizes. A greater proportion of CNVs <5kb are detected in 3kb-mate-pair compared to 5kb-mate-pair WGS (Figure 3b,c). This is likely because for longer insert sizes the experimental variability in size selection during library preparation makes for increased uncertainty in calling smaller CNVs. At the same time, a longer insert size has a greater ability to span CNV boundaries, yielding higher physical coverage, thus increasing overall detection power, especially for CNVs in the medium to large size range. Furthermore, the increases in total CNV detection as a result of increasing coverage are mainly for CNVs <50kb in both read-depth and discordant read-pair analyses (Figure 3d-i).

**Figure 3.**
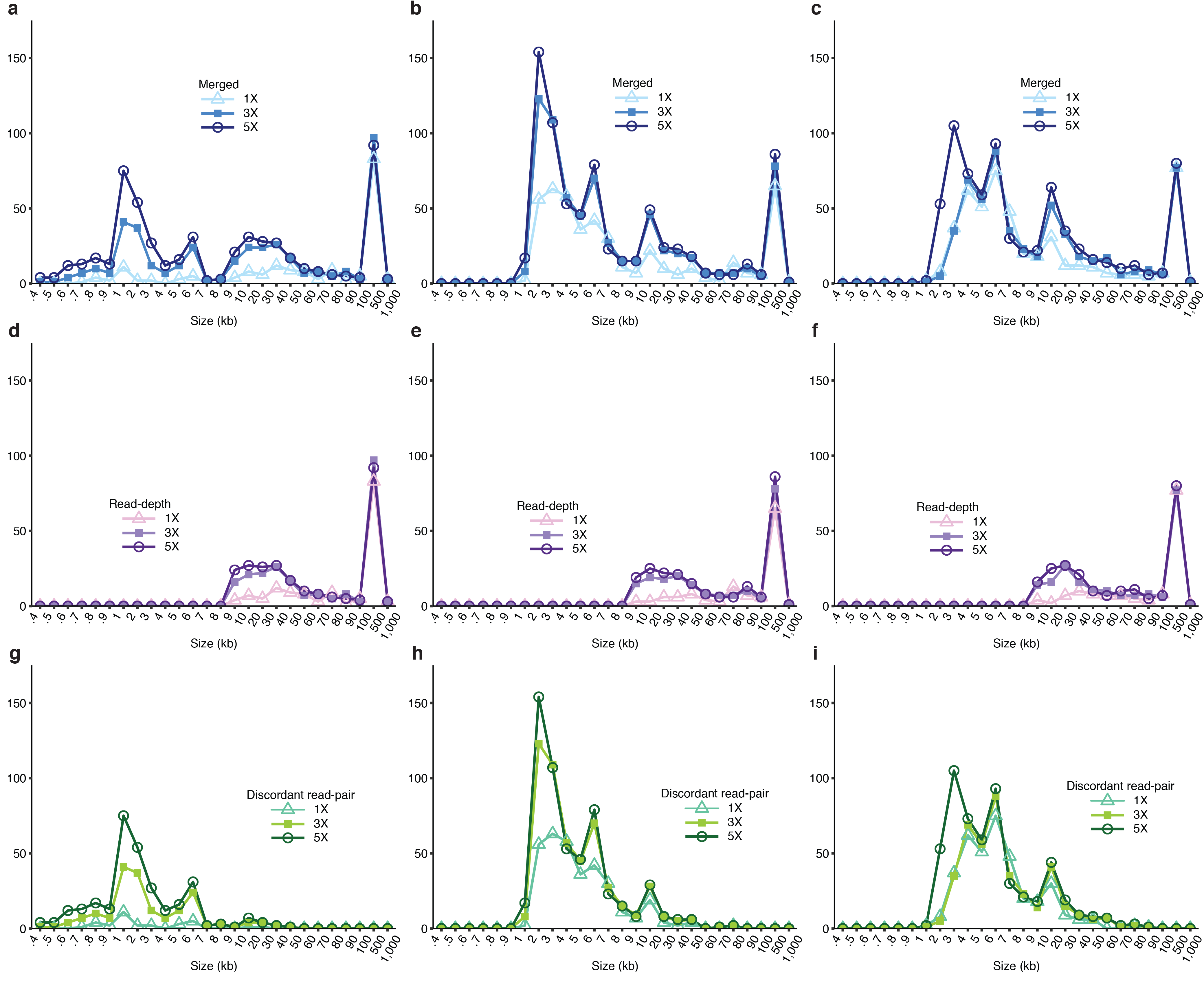
Size Distributions of NA12878 Detected by WGS. Size distribution of NA12878 CNVs detected from (**a**) short-insert, (**b**) 3kb mate-pair, and (**c**) 5kb mate-pair libraries called by combining discordant read-pair and read-depth analysis. CNVs called from (**d**) short-insert, (**e**) 3kb mate-pair, and (**f**) 5kb mate-pair libraries by read-depth analysis only. CNVs called from (**g**) short-insert, (**h**) 3kb mate-pair, and (**i**) 5kb mate-pair libraries by discordant read-pair analyses only.

### All seven GS deletions >100kb are detected by WGS ≥ 1x coverage

All seven GS deletions >100kb are detected with all WGS libraries ≥ 1x coverage. Six deletions are detected with >50% reciprocal overlap, and one deletion (chr6:78,892,808-79,053,430) is detected with a mean overlap = 44% (Supplementary Table S5). Previously, it had been shown that only one of these deletions (chr19:20,595,835-20,717,950) is detected by most arrays [24] (Table 1). Affymetrix SNP 6.0 and Affymetrix CytoScanHD perform the best out of the 17 arrays detecting six and five of these deletions respectively (Table 1). Four out of these seven deletions are detected in Agilent 2x400 CGH but as high copy duplications. The GS deletion on chromosome 3 (chr3: 162,514,471-162,625,647) is only detected in the Agilent CGH arrays but consistently mis-called as a duplication (Supplementary Discussion).

### Detection of the 15 Mbp Cri-du-chat deletion by read-depth analysis

As a vignette with immediate clinical relevance, we also demonstrate that a much larger CNV, the 15 Mbp Cri-du-chat deletion spanning from 5p15.31 to 5p14.2 in NA16595 (sample from Coriell Institute), can be readily detected in a short-insert WGS library at coverages of 1x, 3x, and 5x using read-depth analysis only (Supplementary Table S6). It can also be easily visualized at all coverages by the substantial drop in read-depth coverage in Integrative Genomics Viewer [31] (Figure 4). This NA16595 Cri-du-chat deletion is also confirmed using the Illumina Multi-Ethnic Genotyping Array with two technical replicates (Supplementary Table S6).

**Figure 4.**
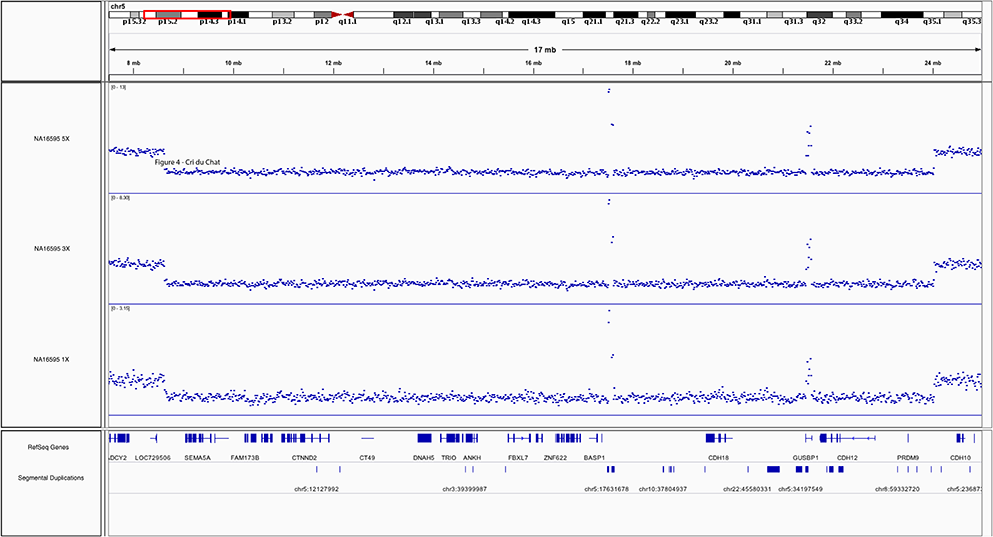
NA16595 Cri-du-Chat Deletion. IGV [31] screenshot of the 15 Mbp Cri-du-Chat deletion on chromosome 5 in NA16595 by short-insert WGS at 1X, 3X, and 5X coverages. Vertical axis: coverage value, blue dots: respective coverage at genomic positions. Two areas within the deletion show unusually high coverage due to overlap with segmental duplications resulting in cross-mapping of sequencing reads.

## DISCUSSION

Using the CNVs in NA12878 as a benchmark, we systematically compared the CNV detection performances of low-coverage WGS strategies relative to each other and relative to various arrays currently in routine-use for cytogenetics. CNVs were called using standard methods for both low-coverage WGS and arrays and then compared to a list of NA12878 GS CNVs that had been distilled from the 1000 Genomes Project [23] as well as to a set of Silver Standard CNVs generated from 1000- Genomes-Project 60x-coverage WGS data (2×250 bp, short-insert) [24]. The Silver Standard CNVs were called using CNVnator [25] which is also used in this study for read-depth analysis of low-coverage WGS data. This further increases the direct comparability of CNV-calling efficiency across the range of coverages though a certain bias in favor of WGS is therefore present in the Silver Standard-based parts of the performance comparison. In almost all scenarios, the WGS approaches show considerably higher sensitivities at detecting GS CNVs than even the best performing arrays (Figure 2a) and are furthermore accompanied by lower percentages of total CNV calls without validation (Supplementary Figure S3, Figure 1b). While all methods of CNV detection left >80% of total GS CNVs undetected, this can be largely explained by that 63% of GS CNVs are <1 kb and 54% are <500 bp which are outside the sensitive detection ranges for all methods (Figure 2b-d, Supplementary Figure S5).

Twenty autosomal GS CNVs are detected collectively in the 17 arrays but not by low-coverage WGS, whereas 426 are detected by low-coverage WGS but not by any of the arrays (Supplementary Table S7). Of these 20 GS CNVs, 4 are approximately 1 kb or shorter, where the sensitivity of detection is low (Figure 2b-d); 13 (65%) are in regions excluded from WGS analysis. These regions include segmental duplications, the MHC cluster, regions that are different between hg19 and hg38 (likely misassembled regions in hg19), and regions in the ENCODE blacklist (regions that often produce artificially high coverage due to excessive unstructured anomalous mapping) [27]. The remaining 3 CNVs do not fall into any of these categories. It is unclear why these 3 CNVs were not detected with WGS, but by visual inspection, their boundaries lie within repetitive elements i.e. LINE1, SINE, segmental duplications. One of these 3 CNVs (chr1: 248756741-248797597) was called as a larger deletion (chr1: 248692001-248820000) in WGS. It is likely that the size this deletion call was extended due to noisy coverage signal from its flanking segmental duplication regions.

Although one may be tempted to conclude that in certain cases arrays-based techniques are superior for CNV detection since there are indeed 20 GS CNVs that are not detected in WGS. However, it is also important to note that these 20 CNVs are detected by combining 17 arrays and that these CNVs are elusive to detection from low-coverage WGS largely due to small size and to occurring in problematic regions excluded from analysis. However, these genomic features are also problematic for arrays in most cases. Our analysis shows that no single array platform or design is specifically sensitive for detecting CNVs that are associated with these features. Therefore, while there are a few specific cases in which CNVs are detected in arrays and not in WGS, we do not see a scenario for which one can make a general statement that array-based techniques are superior for detecting CNVs associated with particular genomic characteristics. In any case, our results show that it is >20 times more likely that a CNV is detected in low-coverage WGS and not in any arrays (Supplementary Table S7)

Although CNV detection methods from WGS data have been available for up to a decade [1, 25, 32, 33], cost, long turnaround times, and heavy computational requirements for deep-coverage WGS analysis have been major obstacles that prevented the adoption of WGS-based methods for cytogenetic applications. Our comparative analysis here shows that these obstacles can be overcome by adopting low-coverage WGS strategies. This means that a cytogenetics laboratory can now avail itself of a technology with a CNV-detection and resolving power that compares very favorably with existing standard methodologies such as arrays or karyotyping while not having to accept an increased burden in terms of cost per sample or turnaround time. Our WGS libraries were sequenced on the Illumina NextSeq 500, in multiplexed fashion, at a cost in sequencing consumables of approximately $150 per 1x coverage. The short-insert libraries were prepared using the Kapa Hyper Prep kit (Kapa Biosystems) where the cost per library is approximately $40-$50 with >50 ng of genomic DNA needed as input for a high-complexity library. The mate-pair library construction reagents cost ~$300 per library using the Illumina Mate Pair Library Prep kit requiring 2 to 4 μg of high molecular weight genomic DNA (mean size >20 kb). The mate-pair library construction costs can decrease further to ~$50-$80 per library if mechanical DNA shearing is employed instead of enzymatic shearing [34]. The costs for arrays are more variable (<$100 for the Illumina PsychArray to several hundred dollars for higher density arrays). Preparation time for arrays (labeling, hybridization, washing, and scanning) and low-coverage short-insert sequencing (library construction, quantification, and loading onto sequencer) both take approximately two days; mate-pair libraries require an additional day. The analysis can all be performed on a standard desktop computer.

The amount of input genomic DNA required for mate-pair WGS is approximately 2-fold more than for most arrays. However, as long as this amount of DNA is available it could be reasoned that it is preferable to use mate-pair WGS for CNV analysis instead of arrays, considering for example that mate-pair WGS even at just 1x coverage is much more sensitive at detecting CNVs than all currently used arrays. For samples with limited DNA, short-insert WGS at just 3x coverage (at a cost of circa $350 per sample using bench-top instruments such as the Illumina NextSeq 500) is still as effective (if not more so in terms of cost and effort) as arrays while easily outperforming arrays in the ability to detect and resolve CNVs. It should also be taken into consideration that sequencing costs will only continue to decrease thus rendering the use of WGS for CNV detection even more cost effective in the foreseeable future.

The choice of algorithm used for CNV analysis is likely to greatly impact the number and accuracy of CNV calls [33, 35]. A comprehensive comparative analysis of these algorithms is beyond the scope of this present study, though work on this matter has been discussed extensively in recent publications [32, 36–38]. For in-depth discussions of CNV analysis tools, approaches, parameters, and challenges as well as performance comparisons with 30x-coverage WGS (Supplementary Figure S6, Supplementary Table S1), see Supplementary Discussion.

Overall, low-coverage WGS approaches are drastically more sensitive at detecting CNVs compared to the best performing arrays (currently commercially available) and are accompanied with smaller percentages of calls without validation.

The prospect of replacing arrays with low-coverage WGS in a cytogenetic context seems promising and essentially at hand. Our results will contribute to the discussion on when and via which route this transition from using arrays to WGS will be plausible in cytogenetics practice.

## CONTRIBUTORSHIP STATEMENT

AEU and BZ conceived and designed the study. BZ and RP performed the experiments. BZ designed the analysis pipeline. SSH, BZ, and XZ performed analysis. RRH contributed code. BZ, SSH, and AEU wrote the manuscript.

## COMPETING INERESTS

The authors declare no competing interest.

## ACKNOWLEDGEMENT

We thank Dr. Billy T. Lau at the Stanford Genome Technology Center for performing the pilot round of library sequencing on the Illumina NextSeq 500.

## References

1 Korbel JO, Urban AE, Affourtit JP, Godwin B, Grubert F, Simons JF, Kim PM, Palejev D, Carriero NJ, Du L, Taillon BE, Chen Z, Tanzer A, Saunders ACE, Chi J, Yang F, Carter NP, Hurles ME, Weissman SM, Harkins TT, Gerstein MB, Egholm M, Snyder M. Paired-end mapping reveals extensive structural variation in the human genome. Science 2007;318:420–6.

2 Frazer KA, Murray SS, Schork NJ, Topol EJ. Human genetic variation and its contribution to complex traits. Nat Rev Genet 2009;10:241–51.

3 1000 Genomes Project Consortium, Abecasis GR, Altshuler D, Auton A, Brooks LD, Durbin RM, Gibbs RA, Hurles ME, McVean GA. A map of human genome variation from population-scale sequencing. Nature 2010;467:1061–73.

4 1000 Genomes Project Consortium, Abecasis GR, Auton A, Brooks LD, DePristo MA, Durbin RM, Handsaker RE, Kang HM, Marth GT, McVean GA. An integrated map of genetic variation from 1,092 human genomes. Nature 2012;491:56–65.

5 1000 Genomes Project Consortium, Auton A, Brooks LD, Durbin RM, Garrison EP, Kang HM, Korbel JO, Marchini JL, McCarthy S, McVean GA, Abecasis GR. A global reference for human genetic variation. Nature 2015;526:68–74.

6 Sebat J, Lakshmi B, Troge J, Alexander J, Young J, Lundin P, Månér S, Massa H, Walker M, Chi M, Navin N, Lucito R, Healy J, Hicks J, Ye K, Reiner A, Gilliam TC, Trask B, Patterson N, Zetterberg A, Wigler M. Large-scale copy number polymorphism in the human genome. Science 2004;305:525–8.

7 de Vries BBA, Pfundt R, Leisink M, Koolen DA, Vissers LELM, Janssen IM, Reijmersdal S van, Nillesen WM, Huys EHLPG, Leeuw N de, Smeets D, Sistermans EA, Feuth T, van Ravenswaaij-Arts CMA, van Kessel AG, Schoenmakers EFPM, Brunner HG, Veltman JA. Diagnostic genome profiling in mental retardation. Am J Hum Genet 2005;77:606–16.

8 Sharp AJ, Hansen S, Selzer RR, Cheng Z, Regan R, Hurst JA, Stewart H, Price SM, Blair E, Hennekam RC, Fitzpatrick CA, Segraves R, Richmond TA, Guiver C, Albertson DG, Pinkel D, Eis PS, Schwartz S, Knight SJL, Eichler EE. Discovery of previously unidentified genomic disorders from the duplication architecture of the human genome. Nat Genet 2006;38:1038–42.

9 Sebat J, Lakshmi B, Malhotra D, Troge J, Lese-Martin C, Walsh T, Yamrom B, Yoon S, Krasnitz A, Kendall J, Leotta A, Pai D, Zhang R, Lee Y-H, Hicks J, Spence SJ, Lee AT, Puura K, Lehtimäki T, Ledbetter D, Gregersen PK, Bregman J, Sutcliffe JS, Jobanputra V, Chung W, Warburton D, King M-C, Skuse D, Geschwind DH, Gilliam TC, Ye K, Wigler M. Strong association of de novo copy number mutations with autism. Science 2007;316:445–9.

10 Walsh T, McClellan JM, McCarthy SE, Addington AM, Pierce SB, Cooper GM, Nord AS, Kusenda M, Malhotra D, Bhandari A, Stray SM, Rippey CF, Roccanova P, Makarov V, Lakshmi B, Findling RL, Sikich L, Stromberg T, Merriman B, Gogtay N, Butler P, Eckstrand K, Noory L, Gochman P, Long R, Chen Z, Davis S, Baker C, Eichler EE, Meltzer PS, Nelson SF, Singleton AB, Lee MK, Rapoport JL, King M-C, Sebat J. Rare Structural Variants Disrupt Multiple Genes in Neurodevelopmental Pathways in Schizophrenia. Science (80- ) 2008;320:539–43.

11 Girirajan S, Campbell CD, Eichler EE. Human Copy Number Variation and Complex Genetic Disease. Annu Rev Genet 2011;45:203–26.

12 Zhang Y, Haraksingh R, Grubert F, Abyzov A, Gerstein M, Weissman S, Urban AE. Child development and structural variation in the human genome. Child Dev 2013;84:34–48.

13 Michels E, De Preter K, Van Roy N, Speleman F. Detection of DNA copy number alterations in cancer by array comparative genomic hybridization. Genet Med 2007;9:574–84.

14 Uddin M, Thiruvahindrapuram B, Walker S, Wang Z, Hu P, Lamoureux S, Wei J, MacDonald JR, Pellecchia G, Lu C, Lionel AC, Gazzellone MJ, McLaughlin JR, Brown C, Andrulis IL, Knight JA, Herbrick J-A, Wintle RF, Ray P, Stavropoulos DJ, Marshall CR, Scherer SW. A high-resolution copy-number variation resource for clinical and population genetics. Genet Med 2015;17:747–52.

15 D’Arrigo S, Gavazzi F, Alfei E, Zuffardi O, Montomoli C, Corso B, Buzzi E, Sciacca FL, Bulgheroni S, Riva D, Pantaleoni C. The Diagnostic Yield of Array Comparative Genomic Hybridization Is High Regardless of Severity of Intellectual Disability/Developmental Delay in Children. J Child Neurol 2016;31:691–9.

16 Stobbe G, Liu Y, Wu R, Hudgings LH, Thompson O, Hisama FM. Diagnostic yield of array comparative genomic hybridization in adults with autism spectrum disorders. Genet Med 2014;16:70–7.

17 Gijsbers AC, Lew JY, Bosch CA, Schuurs-Hoeijmakers JH, van Haeringen A, den Hollander NS, Kant SG, Bijlsma EK, Breuning MH, Bakker E, Ruivenkamp C AL. A new diagnostic workflow for patients with mental retardation and/or multiple congenital abnormalities: test arrays first. Eur J Hum Genet 2009;17:1394–402.

18 Liang D, Peng Y, Lv W, Deng L, Zhang Y, Li H, Yang P, Zhang J, Song Z, Xu G, Cram DS, Wu L. Copy number variation sequencing for comprehensive diagnosis of chromosome disease syndromes. J Mol Diagn 2014;16:519–26.

19 Liu S, Song L, Cram DS, Xiong L, Wang K, Wu R, Liu J, Deng K, Jia B, Zhong M, Yang F. Traditional karyotyping vs copy number variation sequencing for detection of chromosomal abnormalities associated with spontaneous miscarriage. Ultrasound Obstet Gynecol 2015;46:472–7.

20 Dong Z, Zhang J, Hu P, Chen H, Xu J, Tian Q, Meng L, Ye Y, Wang J, Zhang M, Li Y, Wang H, Yu S, Chen F, Xie J, Jiang H, Wang W, Wai Choy K, Xu Z. Low-pass whole-genome sequencing in clinical cytogenetics: a validated approach. Genet Med 2016;18:940–8.

21 Zook JM, Chapman B, Wang J, Mittelman D, Hofmann O, Hide W, Salit M. Integrating human sequence data sets provides a resource of benchmark SNP and indel genotype calls. Nat Biotechnol 2014;32:246–51.

22 Eberle MA, Fritzilas E, Krusche P, Källberg M, Moore BL, Bekritsky MA, Iqbal Z, Chuang H-Y, Humphray SJ, Halpern AL, Kruglyak S, Margulies EH, McVean G, Bentley DR. A reference data set of 5.4 million phased human variants validated by genetic inheritance from sequencing a three-generation 17-member pedigree. Genome Res 2017;27:157–64.

23 Sudmant PH, Rausch T, Gardner EJ, Handsaker RE, Abyzov A, Huddleston J, Zhang Y, Ye K, Jun G, Hsi-Yang Fritz M, Konkel MK, Malhotra A, Stütz AM, Shi X, Paolo Casale F, Chen J, Hormozdiari F, Dayama G, Chen K, Malig M, Chaisson MJP, Walter K, Meiers S, Kashin S, Garrison E, Auton A, Lam HYK, Jasmine Mu X, Alkan C, Antaki D, Bae T, Cerveira E, Chines P, Chong Z, Clarke L, Dal E, Ding L, Emery S, Fan X, Gujral M, Kahveci F, Kidd JM, Kong Y, Lameijer E-W, McCarthy S, Flicek P, Gibbs RA, Marth G, Mason CE, Menelaou A, Muzny DM, Nelson BJ, Noor A, Parrish NF, Pendleton M, Quitadamo A, Raeder B, Schadt EE, Romanovitch M, Schlattl A, Sebra R, Shabalin AA, Untergasser A, Walker JA, Wang M, Yu F, Zhang C, Zhang J, Zheng-Bradley X, Zhou W, Zichner T, Sebat J, Batzer MA, McCarroll SA, Mills RE, Gerstein MB, Bashir A, Stegle O, Devine SE, Lee C, Eichler EE, Korbel JO. An integrated map of structural variation in 2,504 human genomes. Nature 2015;526:75–81.

24 Haraksingh RR, Abyzov A, Urban AE. Comprehensive performance comparison of high-resolution array platforms for genome-wide Copy Number Variation (CNV) analysis in humans. BMC Genomics 2017;18:321.

25 Abyzov A, Urban AE, Snyder M, Gerstein M. CNVnator: An approach to discover, genotype, and characterize typical and atypical CNVs from family and population genome sequencing. Genome Res 2011;21:974–84.

26 Layer RM, Chiang C, Quinlan AR, Hall IM. LUMPY: a probabilistic framework for structural variant discovery. Genome Biol 2014;15:R84.

27 ENCODE Project Consortium. An integrated encyclopedia of DNA elements in the human genome. Nature 2012;489:57–74.

28 McCarroll SA, Kuruvilla FG, Korn JM, Cawley S, Nemesh J, Wysoker A, Shapero MH, de Bakker PIW, Maller JB, Kirby A, Elliott AL, Parkin M, Hubbell E, Webster T, Mei R, Veitch J, Collins PJ, Handsaker R, Lincoln S, Nizzari M, Blume J, Jones KW, Rava R, Daly MJ, Gabriel SB, Altshuler D. Integrated detection and population-genetic analysis of SNPs and copy number variation. Nat Genet 2008;40:1166–74.

29 Kent WJ, Sugnet CW, Furey TS, Roskin KM, Pringle TH, Zahler AM, Haussler D. The human genome browser at UCSC. Genome Res 2002;12:996–1006.

30 Karolchik D, Hinrichs AS, Furey TS, Roskin KM, Sugnet CW, Haussler D, Kent WJ. The UCSC Table Browser data retrieval tool. Nucleic Acids Res 2004;32:D493–6.

31 Robinson JT, Thorvaldsdóttir H, Winckler W, Guttman M, Lander ES, Getz G, Mesirov JP. Integrative genomics viewer. Nat Biotechnol 2011;29:24–6.

32 Zhao M, Wang Q, Wang Q, Jia P, Zhao Z. Computational tools for copy number variation (CNV) detection using next-generation sequencing data: features and perspectives. BMC Bioinformatics 2013;14:S1.

33 Pabinger S, Dander A, Fischer M, Snajder R, Sperk M, Efremova M, Krabichler B, Speicher MR, Zschocke J, Trajanoski Z. A survey of tools for variant analysis of next-generation genome sequencing data. Brief Bioinform 2014;15:256–78.

34 Park N, Shirley L, Gu Y, Keane TM, Swerdlow H, Quail M a. An improved approach to mate-paired library preparation for Illumina sequencing. Methods Next Gener Seq 2013;1:10–20.

35 Tattini L, D’Aurizio R, Magi A. Detection of Genomic Structural Variants from Next-Generation Sequencing Data. Front Bioeng Biotechnol 2015;3. doi:10.3389/fbioe.2015.00092

36 Abel HJ, Duncavage EJ. Detection of structural DNA variation from next generation sequencing data: a review of informatic approaches. Cancer Genet 2013;206:432–40.

37 Lin K, Smit S, Bonnema G, Sanchez-Perez G, de Ridder D. Making the difference: integrating structural variation detection tools. Brief Bioinform 2015;16:852–64.

38 Pirooznia M, Goes FS, Zandi PP. Whole-genome CNV analysis: advances in computational approaches. Front Genet 2015;6. doi:10.3389/fgene.2015.00138

39 Alkan C, Coe BP, Eichler EE. Genome structural variation discovery and genotyping. Nat Rev Genet 2011;12:363–76.

